# Telomere elongation and Telomerase activity in Normal and Cancer cell lines: HEK-293, HeLa and A549

**DOI:** 10.1101/100446

**Authors:** Ankur Patel, Grishma Joshi, Yunke Fu, Sherwin Shaju, Angelo Messina

## Abstract

Telomerase, an eukaryotic ribonucleoprotein (RNP) complex, consists of an essential RNA template and a reverse transcriptase termed as human telomerase reverse transcriptase (hTERT). By using this reverse transcriptase enzyme, telomerase RNP maintains its telomeric length in all cancer cells and in few stem cells. As cells age, telomerase level decreases and telomere get shorten, leading to senescence. Presence of telomerase helps delay the senescence due to active production of telomere, leading to cell turning malignant and eventually cancerous. In this experiment, we would investigate the telomerase activity tested using q PCR and telomerase level using western blot in three different cell lines, HEK-293, HeLa and A549.

**Hypothesis:** Where HEK-293 cells have longer telomeres and higher telomerase activity, HeLa and A549 cell lines will have varied length of telomere but higher telomerase activity.

## Introduction

A principal distinction in the conduct of normal cells versus tumor cells in culture [1–5] is that normal cells replicate for a set number of times (display cell senescence) while tumor cells for the most part can multiply uncertainly and are often termed as immortal [6]. The essential part of human cells that influence cell aging are telomeres. As a result of DNA replication process, each time the chromosomes are replicated, the ends get shorter. This chromosomal breakdown would soon prompt to the loss of critical DNA and soon thereafter a cell could not replicate anymore. A telomere is a distinct region of nucleotide sequences at every end of a chromosome, which shields the end of the chromosome from decay or from combination with neighboring chromosomes [7]. Telomeres are G rich T2AG3 repeat sequences at the end of the chromosomes, which are crucial for the survival of tumor cells. As an everyday cell division process, a small piece of telomeric DNA is lost with each cell division. When telomere length reaches a vital limit, the cell undergoes senescence. Telomere length may also therefore serve as a biological clock to decide the lifespan of a cell and an organism. [8] These telomeres are maintained by an enzyme, Telomerase acting as a reverse transcriptase in the vast majority of Tumor cells [9]. The protein component of telomerase, Telomerase Reverse Transcriptase (TERT), is an enzyme. Also, its amino acid sequence includes reverse transcriptase motifs [10]. It is in that context that we are keen on interceding in the telomerase activity. In this experiment, we highlight factors that may antagonistically influence health and lifespan of a person by telomere shortening. According to the hypothesis stated above, the experiment should elucidate the telomeric length of each of the given two cell lines, A549 and HeLa, with longer telomere then HEK-293.

## Material and Methodology

### Cell Line Samples

Three Different types of cell lines were used for investigating the relationship between telomerase and cancer. Cell lines used, HEK-293, HeLa and A549 were also of interest as to investigate how the telomerase differ in each of the cell lines. HEK-293 cells are Human Embryonic Kidney cells. HeLa cells are derived from cervical carcinoma cells and A549 cells are derived from a lung carcinoma.

### TRAP Assay/Bradford Assay

Since, telomerase synthesis telomeres, determination of telomeric DNA synthesized on an oligo template in twenty minutes was performed quantitatively. The assay used is called as TRAP assay (Telomeric Repeat Amplification Protocol) [11]. In this assay, telomerase from the cell extracts of the given cell lines is given twenty minutes to synthesize telomeres on the TS oligo template. The synthesis products are then detected by a variation of q PCR using the same TS oligo template as a forward primer and an Amplifluor reverse primer that binds to the telomere repeat. The amount of telomere produced by the telomerase in the cell extracts by measuring the production of fluorescence during PCR.

q PCR program.

q PCR program,
| <u>Step</u> | <u>Time</u> | <u>Temperature</u><br>(°C) | <u>Comments</u> |
| --- | --- | --- | --- |
| Telomerase reaction | 30 min | 30 | Telomere templates formed by reaction adding 6-base-repeats to primer with the activity of telomerase HotActivitedTaq DNA Polymerase Activation Step is activated by this heating step. |
| PCR initial activation | 2 min | 94 |  |
| 3 Step cycling 45 times: |  |  |  |
| Denaturation | 30 sec | 94 |  |
| Annealing | 30 sec | 59 |  |
| Extension | 1 min | 72 |  |
| Final Extension | 3 min | 72 |  |

### Polyacrylamide Gel

The q PCR results tell us which cell extracts produced more telomeric DNA. A more active telomerase could synthesize longer telomeres, synthesizes telomeres on more TS oligos, or both. To differentiate these possibilities, q PCR products are run on Polyacrylamide gel (PAGE). If the telomerase synthesizes longer telomeres, larger bands than a telomerase that synthesized telomeres on more TS oligos should be observed.

### Western Blot

Variation in cellular telomerase activity could result from two major differences between cell lines: a difference in the amount of telomerase in the cell or a change in the telomerase that allows it to more efficiently synthesize telomeres. To test that the differences in telomerase activity results from difference in telomerase levels, Western blots were performed on the same cell extracts used earlier. Telomerase (~130kDa) was probed over night with α-telomerase antibody diluted 1:500, to see if the cells with greater telomerase activity have greater amounts of telomerase protein in the cells. As a control, GAPDH (~40kDa) was probed overnight for housekeeping gene with α- GAPDH antibody dilutes 1:10,000.

### Observation and Results

The absorbance for the Bradford assay were measured for the standards *(Figure 1),* using which we calculated the amount of protein(telomerase) in each given cell lines, i.e. HEK-293= 0.583 μg/μl, A549= 12.82 μg/μl and HeLa= 10.0 μg/μl. The synthesis products are then detected by a variation of q PCR.

**Figure 1.**
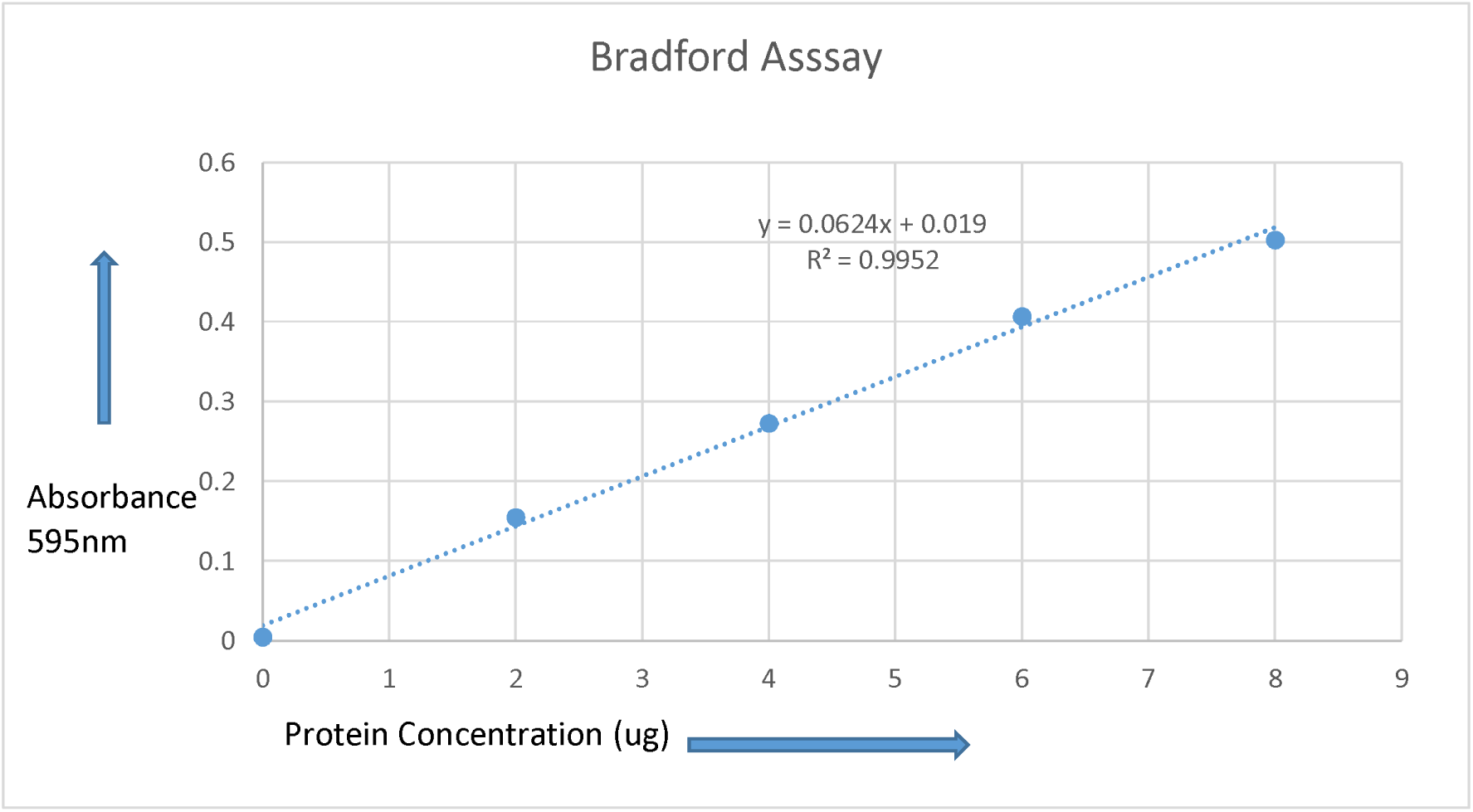
Bradford Assay. *The absorbance at 595nm was measure for samples containing 2,4,6 and 8 μg BSA. The equation can be used to determine the concentration of the Cell Lines.*

Using the q PCR raw data, graphical presentation was then performed, in which cell lines #4,5 and 6 showed less fluorescence activity whereas, cell lines #7,8 and 9 were observed with much higher fluorescence. *(Figure 2).* Also to determine how much telomerase was present per reaction time, cycle threshold standard graph was plotted.

**Figure 2.**
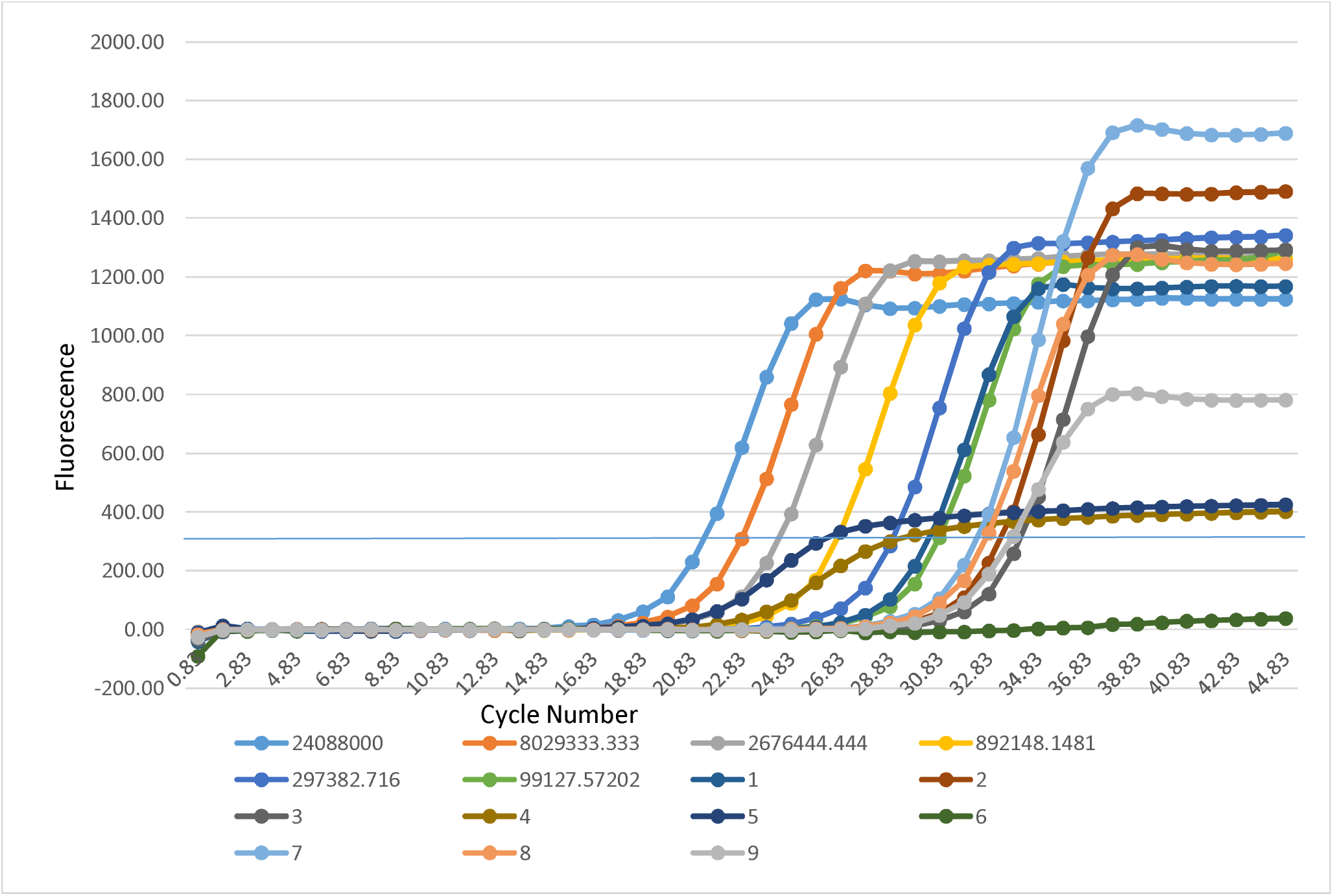
Graph for qPCR results. *SYBR green real time PCR was performed on TSR standards of the given concentrations and telomere synthesis reactions using native or heat treated cell extracts from given Cell lines. The black line at ~300 represent cycle at which each PCR crossed the fluroscence at ~300 nm.*

**Figure 3:**
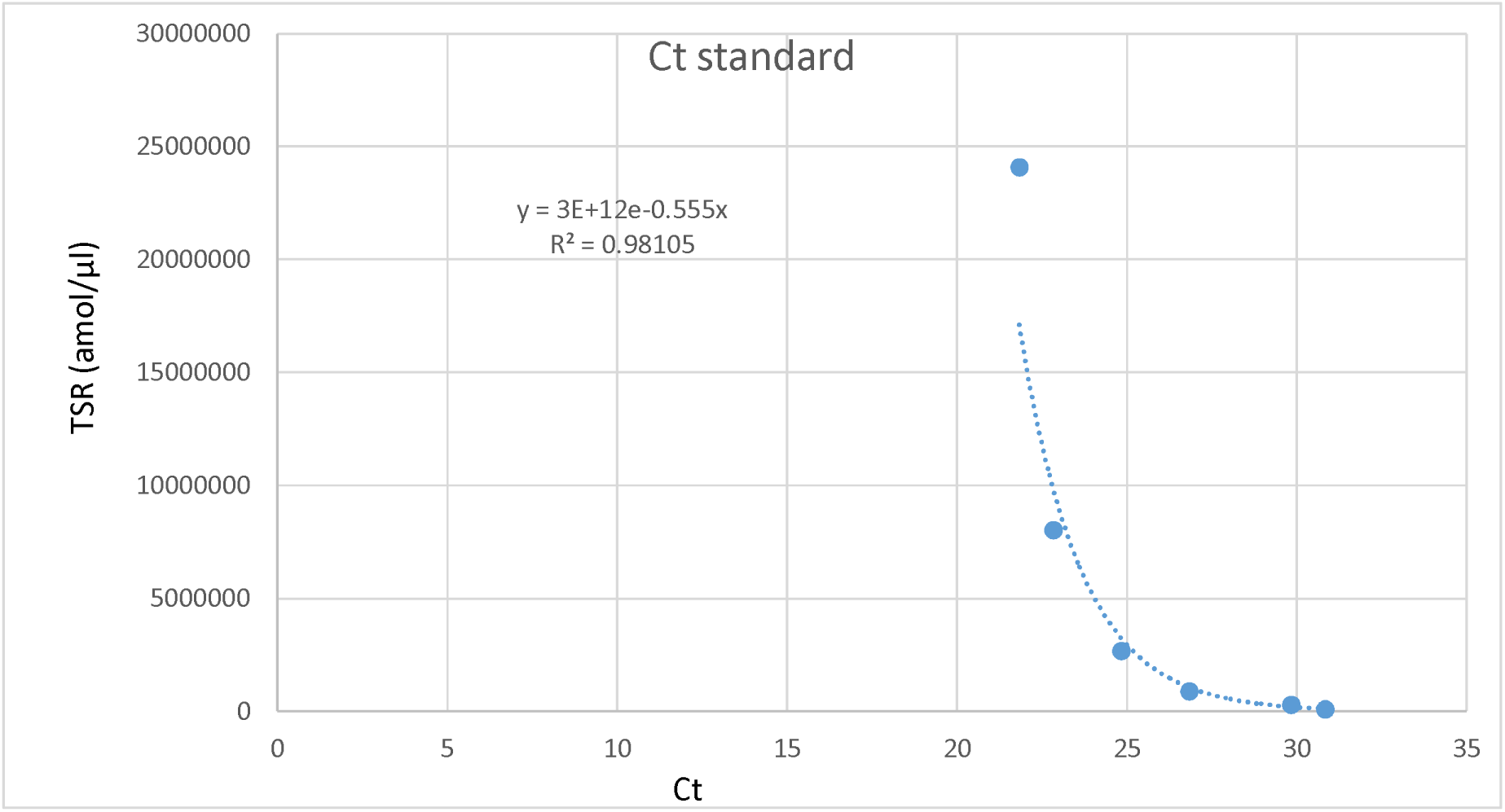
Cycle threshold Standard. Using which teloemrase per reaaction would be calculated.

| Keys for Figure 2. |  |
| --- | --- |
| 1 | Positive control |
| 2 | Lysis buffer |
| 3 | PCR H<sub>2</sub>O |
| 4 | HEK-293 (85 °C) |
| 5 | A549 (85 °C) |
| 6 | HeLa (85 °C) |
| 7 | HEK-293 |
| 8 | A549 |
| 9 | HeLa |
| Ct | Cycle Threshold |

Cycle threshold standard graph was performed using which Telomerase per reaction time was calculated [Table 1.]

**Table 1:**
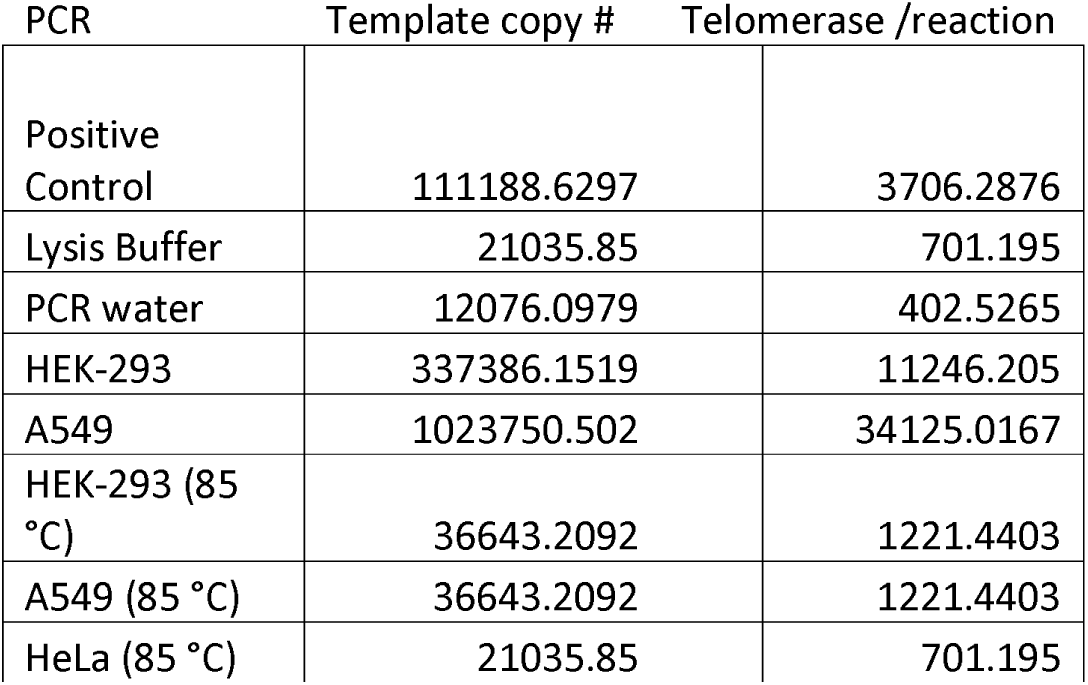
*Amount of telomerase per reaction time*

| PCR | Template copy # | Telomerase /reaction |
| --- | --- | --- |
| Positive Control | 111188.6297 | 3706.2876 |
| Lysis Buffer | 21035.85 | 701.195 |
| PCR water | 12076.0979 | 402.5265 |
| HEK-293 | 337386.1519 | 11246.205 |
| A549 | 1023750.502 | 34125.0167 |
| HEK-293 (85 °C) | 36643.2092 | 1221.4403 |
| A549 (85 °C) | 36643.2092 | 1221.4403 |
| HeLa (85 °C) | 21035.85 | 701.195 |

Telomerase was observed in PCR (Table 1), except in native HeLa cell lines. To determine whether telomerase is used to either produce long telomeric repeats on few templates or short repeats are produced using more templates, q PCR products were ran on PAGE.

Telomeres were observed with higher intensity in the native cell extracts rather than heat killed cell extracts, and were found at the bottom part of the gel. Also telomere banding pattern were observed in all other PCR samples (Figure 4). To find out the telomerase intensity in the samples, western blot was performed using GAPDH as a control housekeeping gene.

**Figure 4:**
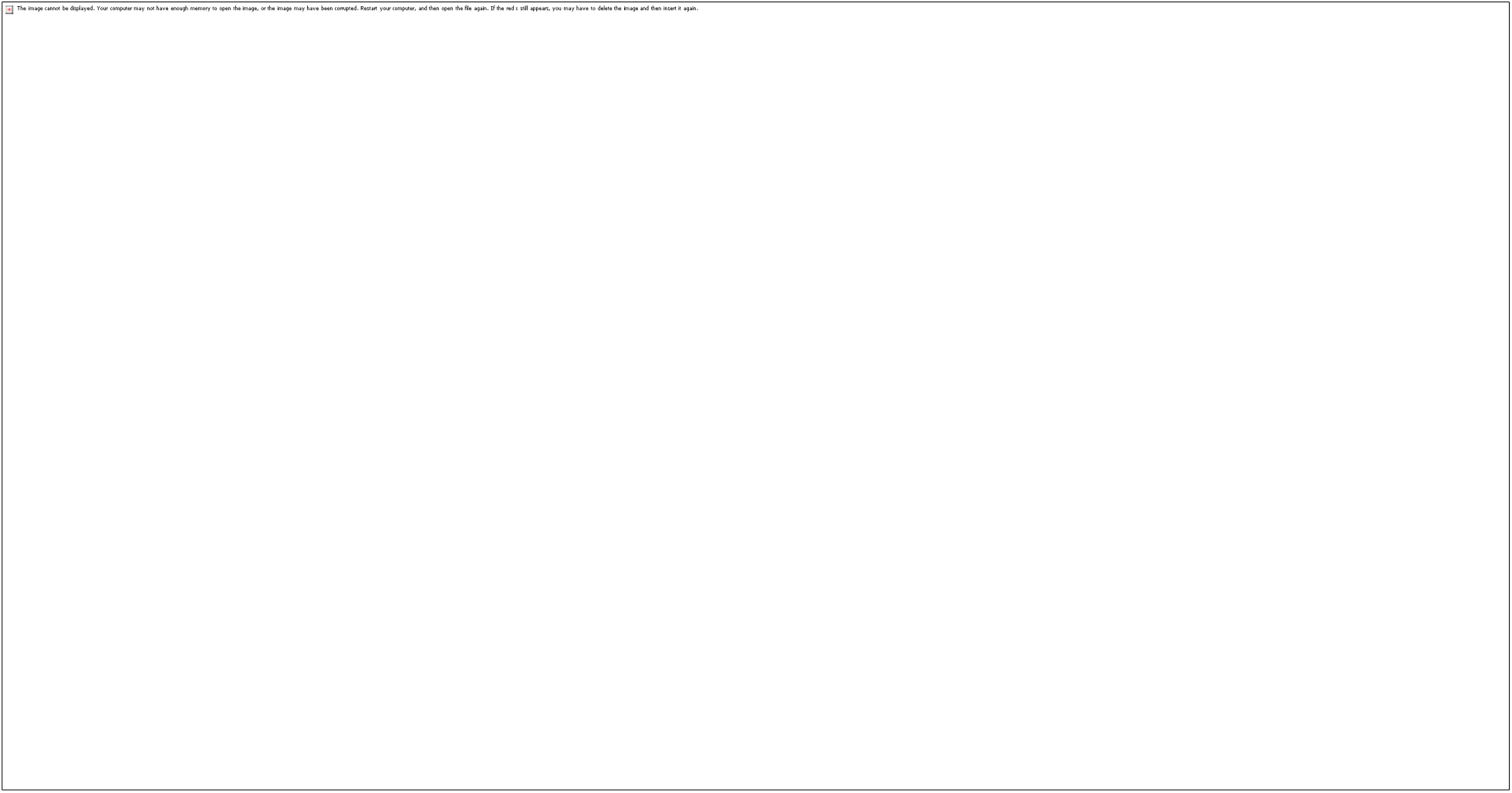
POLYACRYLAMIDE GEL ELECTROPHORESIS, for detection of telomere length in each given cell lines.

**Figure 5:**
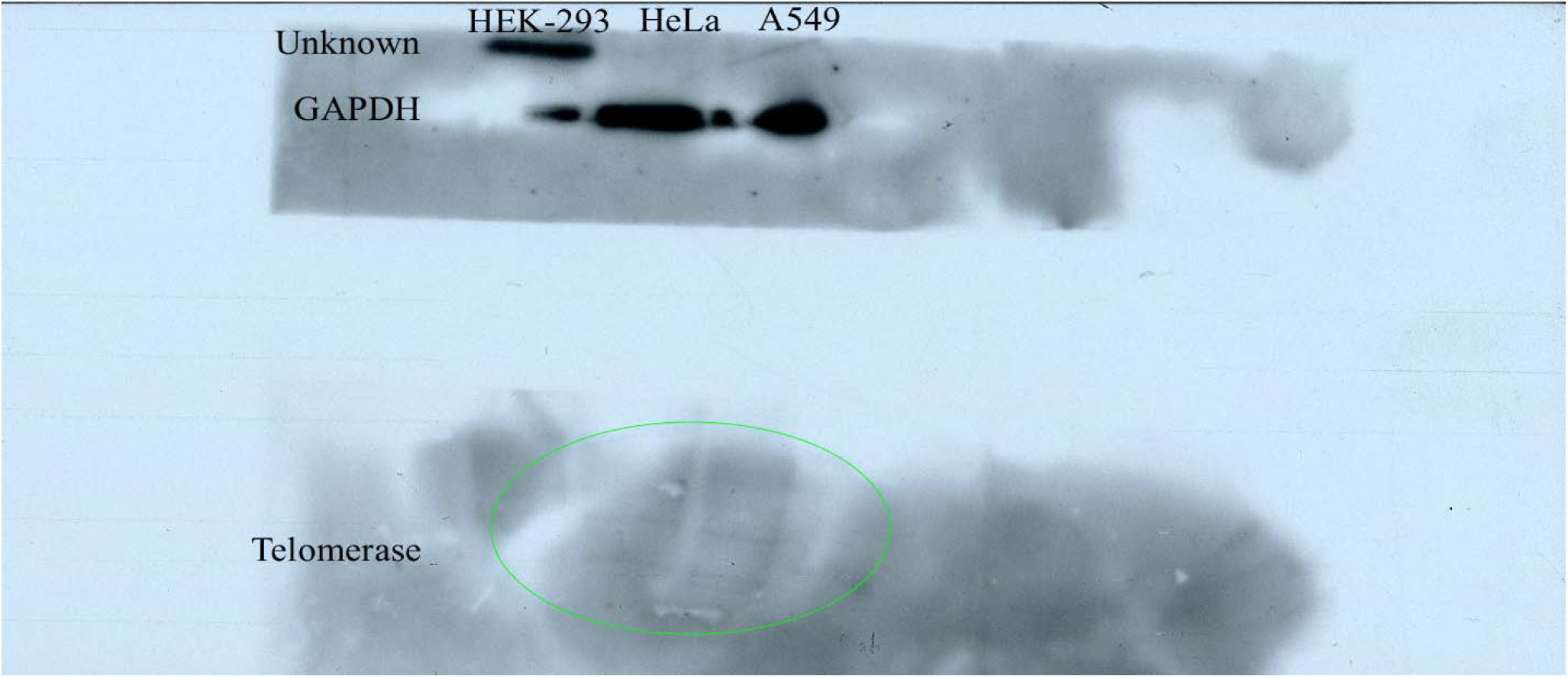
Western Blot for telomerase and GAPDH.

Banding pattern in the western blot was observed, with faint telomerase bands in HeLa and A549 but fainter in HEK-293.

## Conclusion and Discussion

In the q PCR, since, the series 1, shows higher telomerase activity as it is a positive control, but within series 7, 8 and 9, native HeLa (series 9) showed the least activity then HEK-293 (series 7) and A549(series 8). Also the, heat killed cell lines extracts showed very low telomerase activity, as during heat killing process the telomeric DNA was denatured and what got amplified was the genomic DNA. Graph also represents telomerase activity with series 2 (Lysis buffer) and 3 (PCR water), which indicates cross contamination.

Since, the banding pattern in all the PCR series were found at the bottom part of the PAGE gel, it suggests that the telomere produced were shorter repeats with more templates, also the band intensity states that native cell line extracts showed much more intense banding pattern then the other PCR samples. Also, the faint bands in the series 4 at the very bottom of the gel might have occurred due to production of primer dimer.

To measure telomerase intensity, western blots was then performed. Western blots showed faint banding pattern in which HEK-293 showed the least banding pattern. It might be due to the result that over a course of time HEK-293 decreased some amount of telomerase in it. Also, comparatively faint bands from A549 and HeLa cell extracts indicates that there is more telomerase in these cancer cell lines, which leads it to a more cell life in cancer cells, by producing more telomere repeats.

There are some limitation to this experiment, e.g. false negative results might be due to the mutation occurred in the telomeric DNA, which can be detected using sequencing.

## Acknowledgement

Author(s), would like to thank, Shaju Sherwin and Messina Angelo, for their help in preparation of the Lab protocol requirements.

